# Single-Nuclei RNA Sequencing Identifies Type C Low-Threshold Mechanoreceptors as Key Players in Paclitaxel-Induced Peripheral Neuropathy

**DOI:** 10.1101/2023.12.31.573703

**Authors:** Wuping Sun, Rongzhen Li, Xinyi Zhang, Songbin Wu, Yanjun Jiang, Qian Li, Di Cao, Donglin Xiong, Lizu Xiao, Xiaodong Liu

## Abstract

Neuropathic pain triggered by chemotherapy poses a significant clinical challenge. Investigating cell type-specific alterations through single-cell transcriptome analysis holds promise in understanding symptom development and pathogenesis. In this study, we performed single nuclei RNA (snRNA) sequencing of dorsal root ganglions (DRG) to explore the molecular mechanism underlying paclitaxel-induced neuropathic pain. Mouse exposed to repeated paclitaxel doses developed persistent pain hypersensitivity lasting at least 21 days. The snRNA sequencing unveiled seven major cell types within DRGs, with neurons further subdivided into 12 distinct subclusters using known markers. Notably, type C low-threshold mechanoreceptors (C_LTMR) exhibited the most pronounced transcriptomic changes post-paclitaxel administration. Differential gene expression and Gene Ontology (GO) analysis highlighted suppressed potassium-related currents, microtubule transport, and mitochondrial functions in C_LTMR following paclitaxel treatment. Meanwhile, Gene Set Enrichment Analysis (GSEA) suggested increased Interleukin 17 production in C_LTMR after paclitaxel exposure. Pseudo-time analysis uncovered nine distinct states (state 1 to 9) of C_LTMR. State 1 exhibits higher prevalence in paclitaxel-treated mice and altered neurotransmission properties, likely contributing to paclitaxel-induced pain hypersensitivity. This comprehensive exploration sheds light on the molecular mechanisms driving paclitaxel-induced neuropathic pain, offering potential avenues for therapeutic intervention.

## Introduction

Neuropathic pain commonly emerges as an unfavorable consequence of anticancer therapies utilizing various chemotherapeutic agents such as paclitaxel, oxaliplatin, and vincristine. This type of pain is typically manifested by neuropathic symptoms such as numbness and abnormal sensations in the extremities, particularly increased pain perception to mechanical and temperature stimuli, and is therefore referred to as chemotherapy-induced neuropathic pain (CINP) [1, 2]. CINP impacts approximately 30-40% of patients receiving cancer treatments, often leading to an unintended decrease in therapeutic dosage or even discontinuation of treatment [3, 4]. In some cases, CINP can endure for months to years after treatment completion, severely impacting patients’ quality of life [3]. Effective prevention and treatment of CIPN stands as a medical necessity.

Currently, there are no effective clinical strategies to counteract CIPN [5, 6]. The obstacles in developing CIPN-related drugs have several reasons. First, the mechanisms of CIPN are intricate and not entirely understood. Chemotherapy treatments induce multifaceted structural and functional changes in non-cancerous cells, resulting in damage to cell organelles (e.g. cytoskeleton and mitochondrion), altered signaling pathways, changes in membrane receptors and ion channels, and modulation of neurotransmitter levels. These alterations impact the RNA and protein profile of both primary sensory neurons and glial cells and influence the excitability and firing threshold of primary sensory neurons [7–12], contributing to the development of CIPN. In our prior study, we conducted transcriptome analysis of dorsal root ganglions (DRG) in oxaliplatin- and paclitaxel-induced CIPN. We revealed substantial distinctions in the pathogenesis of these two CIPN models. Specifically, paclitaxel tended to directly impact somatosensory neurons, while oxaliplatin exhibited a propensity to affect glial cells [13, 14]. Second, the presence of cancers, which might share common or similar signaling pathways but have opposing effects on CIPN, adds to the complexity of the pathogenesis of CIPN [12]. Preventing CIPN without compromising chemotherapy efficacy presents significant operational challenges. Understanding CIPN-specific mechanisms is pivotal for developing clinical management strategies, holding significant scientific and clinical translational value.

Recent advancements in omics at a single-cell resolution not only elucidate the cellular atlas of pain pathways but also unveil cell-type-specific alterations. These insights enhance our comprehension of symptom emergence, such as tactile allodynia and thermal pain, and reveal dysregulated genes that might have been overlooked in bulk tissue-based analyses [15, 16]. These discoveries undoubtedly introduce a novel dimension to our understanding of the pathogenesis of CIPN and the identification of targets/signaling pathways specific to subtypes of somatosensory neurons. This advancement has the potential to significantly enhance the effectiveness of symptom management or CINP prevention while maintaining the efficacy of anti-cancer therapies. In this study, we aim to deepen the understanding of paclitaxel-induced neuropathic pain by employing single-nuclei RNA sequencing technology. This study may contribute to the development of mechanism-oriented approaches for treating CIPN.

## Materials and methods

### Animals

C57BL/6j mice weighing between 20 and 22 g were provided by the Guangdong Province Laboratory Animal Center (Guangzhou, China), maintained and bred in a normal 12 h light/dark cycle with standard feed and water. Experiments were performed using 8-to 12-week-old male littermates. All experimental procedures were approved by the Animal Care and Use Committee of the 6th Affiliated Hospital of Shenzhen University Medical School in advance and were strictly in compliant with the guideline for the care and use of laboratory animals.

### CIPN model

Paclitaxel was dissolved in a solvent containing mix of DMSO-PEG300-Tween 80-Saline (2:4:1:13). Each mouse received 4 mg/kg of paclitaxel every other day, for a cumulative intraperitoneal dosage of 32 mg/kg, or vehicle control. Intrathecal injections of virus in total volume of 5 µl volume or PBS as vehicle control was administered using a 30-gauge (0.5’) needle and Hamilton syringe under isoflurane anesthesia. Tail flicks were observed in mice as evidence of successful access to the dural cavity for intrathecal injection [17].

### Behavioral analysis

#### Mechanical allodynia (electronic von Frey test)

Mechanical paw withdrawal threshold was assessed with eight von Frey hairs of different bending forces (0.008, 0.02, 0.04, 0.07, 0.16, 0.4, 0.6, and 1.0 mg) as we previously reported [18]. A positive response consisted of flexion, leg raising, or foot licking behavior upon applying the filament to the hind paw. Mice were placed individually in a clear acrylic behavioral chamber. After habituated at least 20 min, von Frey hair was applied to the lateral plantar side of the right hindpaw for 3-5 s to measure the mechanical threshold. Each filament was applied at most 5 times, switched to the next filament of either smaller force (if positive response occurred 3 times), or bigger force (if negative response occurred 3 times).

### Thermal/cold hyperalgesia (hot/cold-plate test)

Thermal hyperalgesia and clod allodynia were assessed in C57BL/6 using a hot/cold plate analgesia meter (BIO-CHP, Bioseb, France) as we previously reported [19]. The hot and cold plates were maintained at 53 ± 1°C and 4 ± 1°C, respectively. The time taken from the moment that exposing the mid-plantar surface of the hind paw on the hot/cold plate to the first nociceptive response including paw licking, lifting, or jumping was regarded as the thermal/cold withdrawal latency and recorded in seconds. A cut off time of 20 s was set to avoid tissue damage. A maximum stimulation time of 30 s was adopted to avoid tissue injury. Each mouse was measured 3 times with a 10-minute interval.

### Nuclei isolation

Nuclei were isolated from mouse dorsal root ganglia (DRGs) as described previously with minor modification [20]. DRGs were dissected in a 1.5 ml Eppendorf tube containing homogenization buffer (20 mM Tris-HCl Ph 8.0, 500 mM sucrose, 50 mM KCl, 10 mM MgCl_2_, 0.1 mM DTT, 1% BSA, 0.1% NP40) supplemented with 1xprotease inhibitor cocktail (Roche, Indianapolis, USA) and 0.4 U/μl RNase inhibitor (Invitrogen, Carlsbad, USA). Tissues were transferred into a 2 ml Dounce Tissue Grinder (Kimble Chase, Rockwood, USA) and homogenized with A (“loose”) pestle for 20 strokes, followed with 20 strokes using B (“loose”) pestle on ice. The solution was filtered using a 70-µm cell strainer (Miltenyi Biotec, Maryland, USA) into a 15 ml tube, then filtered with a 30-µm cell strainer (Miltenyi Biotec, Maryland, USA). Nuclei were pelleted with centrifugation at 300 g for 5 min at 4 °C and resuspended in 1.5 ml PBS supplemented with 1% BSA and 0.2 U/μl RNase inhibitor (Invitrogen, Carlsbad, USA). The nuclear suspension was centrifuged at 300 g for 5 min at 4 °C and the pellet was resuspended with 1.5 ml cold PBS.

### Single-nucleus RNA-seq library preparation and sequencing

The nuclei were loaded into microfluidic chip of Chip A Single Cell Kit v2.0 (MobiDrop, Hangzhou, China) to generate droplets with MobiNova-100 (MobiDrop, Hangzhou, China). Each nuclear was wrapped into a droplet which contained reaction reagent and a gel bead linked with up to millions oligos (cell unique barcode). After encapsulation, droplets suffer light cut by MobiNovaSP-100(MobiDrop, Hangzhou, China) following oligos diffusion into reaction mix. The mRNAs were captured by gel beads with cDNA amplification in droplets. Following reverse transcription, cDNAs with barcodes were amplified, and a library was constructed using the High Throughput Single Cell 3’RNA-Seq Kit v2.0 (MobiDrop, Hangzhou, China) and the 3’ Single Index Kit (MobiDrop, Hangzhou, China). The snRNA libraries were sequenced on GenoLab M platform (GeneMind Biosciences, ShenZhen, China) via PE150 model [21].

### Statistical analysis

For behavioral tests, the data were analyzed in R and reported as Mean±SEM. For gene expression, the results were presented in box and whisker plots. The middle line in the box represented the median of the data. The bottom and top of the box were the first (Q1) and third (Q3) quartiles, respectively. The whiskers extended from Q1 and Q3 to the most extreme data points, excluding outliers. Outliers were defined as data points less than or greater than 1.5*IQR (inter-quartile range) of Q1 or Q3 and displayed as black dots. Data were tested for assumptions of normality (Shapiro-wilk normality test) and equal variance (Bartlett test of homogeneity of variance) before comparisons of means. When the data were normally distributed and had equal variance between the groups, a parametric one-way or two-way repeated-measures ANOVA followed by multiple comparisons corrected by Tukey’s Honest Significant Difference (HSD) was applied, otherwise a non-parametric ANOVA (Kruskal-Wallis test with Conover-Iman test to correct multiple comparisons) was used. An adjusted *p*<0.05 was considered a statistically significant difference.

## Results

### Repeated administration of paclitaxel resulted in pain hypersensitivity in mice

We established a mouse model of paclitaxel-induced neuropathic pain according to the outlined procedure in Figure 1A. We assessed mechanical allodynia, as well as thermal and cold hyperalgesia post-paclitaxel administration. As expected, a significant decrease in mechanical threshold was observed on day 3 post-injection, reaching the lowest levels on day 14 post-injection (Figure 1B). Meanwhile, paw withdrawal latencies in response to thermal or cold stimulation declined on day 5 post-injection and were most pronouncedly affected on day 14 post-injection (Figure 1C and D). These observations indicate that the repetitive administration of paclitaxel induced allodynia and hyperalgesia in mice.

**Figure 1.**
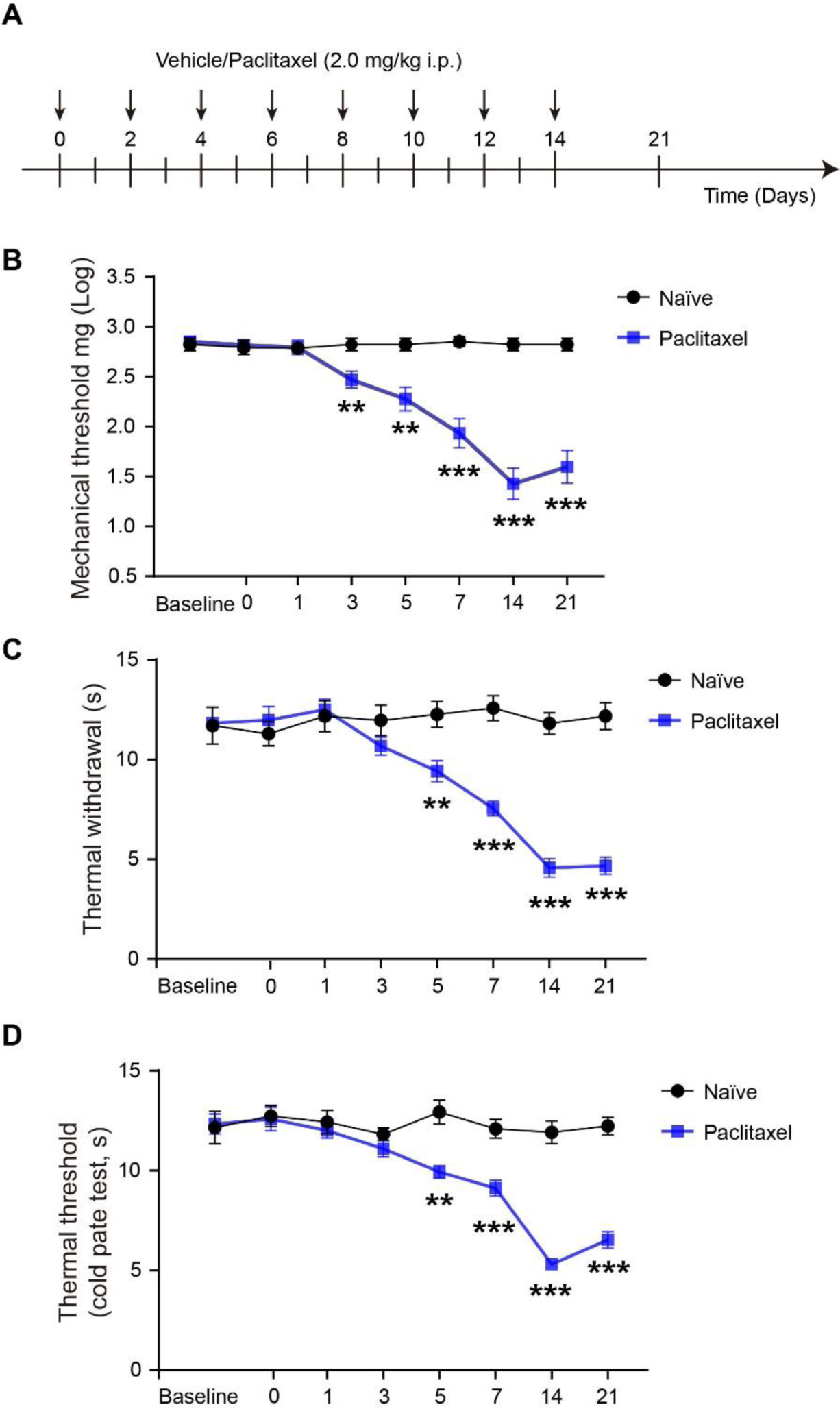
The development of a mouse model of paclitaxel-induced CIPN. **A** The process outline of development of a mouse model of paclitaxel-induced CIPN. **B, C, and D** Mechanical allodynia (B), and thermal (C) and cold hyperalgesia (D) in mice following repeat administration of paclitaxel. The paw withdrawal thresholds in the von Frey test and response latencies in the hot and cold plate tests were measured before treatment (baseline) and after the first injection of either the vehicle or paclitaxel. The bar represents Mean ± SEM, with n=6. The statistical significance is denoted as \*\**p* < 0.01, \*\*\**p* < 0.001 vs. the naive group, and the statistical analysis was performed using two-way repeated-measures ANOVA with Tukey’s HSD correction.

### The neuronal and non-neuronal composition of DRG

To investigate cell type-specific pathogenesis in mice with paclitaxel-induced neuropathic pain, we conducted a single-nuclei RNA sequencing on lumbar (L) 4, 5, and 6 DRGs. The DRGs were isolated from mice (n=x/group) subjected to treatments involving saline for 14 days (referred to Ctrl), paclitaxel treatment for 14 days (referred to Pac14) and paclitaxel treatment followed by a recovery period of 7 days (referred to Pac21). The Fastq files were pre-processed and then mapped with the mouse mm10 reference genome to create expression matrix, cells and genes documents using MobiVision V3 software (MobiDrop). The processed data were loaded separately to the Seurat package (V5) and merged without integration. We filtered out cells exhibited mitochondrial genes percentage exceeding 5%, RNA counts fewer than 500, RNA counts over 5-fold of the median RNA counts, and number of features exceeding 5-fold of the median RNA features. The merged data underwent analysis using “LogNormalize” normalization and “pca” dimension reduction. Subsequently, the first 30 dimensions were input for constructing SNN group, with a resolution of 0.1 applied to differentiate various cell clusters. A total of 8 major cell populations were identified in the DRG (Figure 2A), including Snap25+ neurons, Apoe+ satellite glial cells (SGC), Mpz+ Schwann cells (Schwann), Dcn+ Fibroblasts (Fibro), Flt1+ endothelia cells (Endo), Myl1+ vascular smooth muscle cells (VSMC), and Ptprc+ immune cells (Immune) (Figure 2B-2F). One cluster could not be defined and categorized as undetermined (Und, possibly contaminated cells) due to a lack of known cell type-specific markers (Figure 2B).

**Figure 2.**
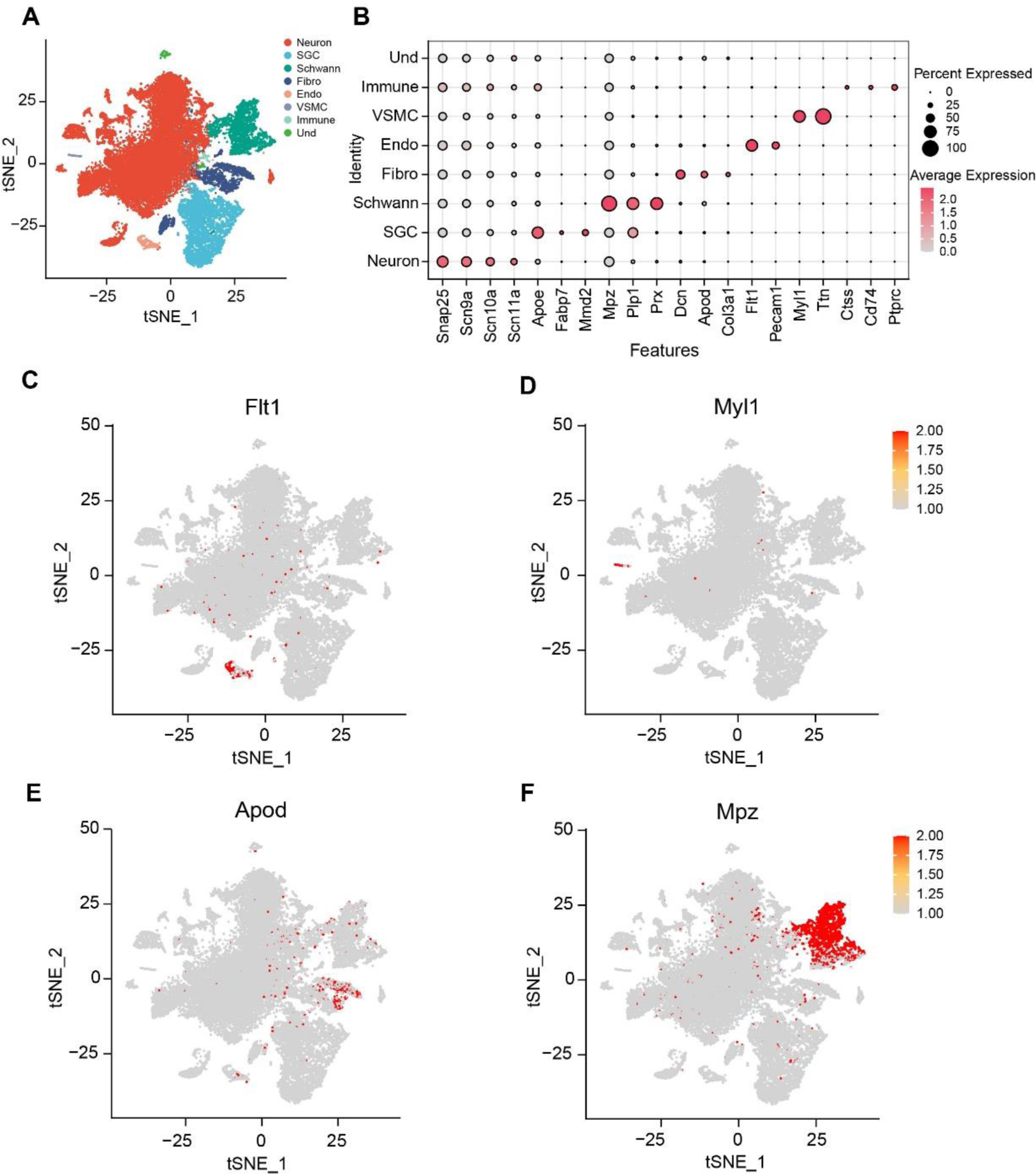
The heterogeneity of dorsal root ganglia (DRG) cells in mice with paclitaxel-induced CIPN. **A** t-distributed scholastic neighbor embedding (t-SNE) plot of single cells profiled in the study, with each cell type represented by a colored dot. **B** The dot plot of the cell types in the DRG of paclitaxel-induced CIPN mice, marked by distinct genes. **C, D, E, and F** t-SNE plots of single cell types of Endo (C), VSMC (D), Fibro (E), and Schwann cells (F), respectively.

### The somatosensory neuron clusters in DRG

We then focused on the neuron cluster and utilized the list of DRG neuron markers summarized by Wang et al. [15] to discern distinct neuronal populations within the DRG. We subset the counts of neuron cluster and perform a similar analysis as described above. The first 30 dimensions were used in PCA reduction and a resolution of 0.5 was set to distinguish neuronal subclusters. We noticed that two clusters (cluster 0 and 1) had very low RNA counts (counts of all cells < 1000) and lacked specific markers. These cells were considered low quality and removed by setting the threshold at nCount_RNA >1000. The remaining data were then re-analyzed. We successfully identified 12 neuronal clusters, including Wnt7a+ proprioceptors (Proprio/Wnt7a), Prokr2+ proprioceptors (Proprio/Prokr2), Ptgfr+ Aβ low-threshold mechanoreceptors (Aβ_LTMR), Cadps2+ Aẟ_LTMR, Tafa4+ C_LTMR, Th+/Tafa4+ C_LTMR/Th, Sstr2+ thermal nociceptors (Heat), Rxfp1+ thermal nociceptors (Heat/Rxfp1), Trpm8+ cold nociceptors (Cold), Mrgprd+/Lpar3+ pruriceptors (Itch/Lpar3), Nppb+ pruriceptors (Itch/Nppb) and Mrgpra3+ pruriceptors (Itch/Mrgpra3). A small cluster included multiple markers and was considered a mixed cell cluster. The t-SNE and UMAP coordinates of all neurons were plotted and demonstrated (Figure 3A-3B). The dot plot of neuronal subtype-specific markers was also displayed (Figure 3C).

**Figure 3.**
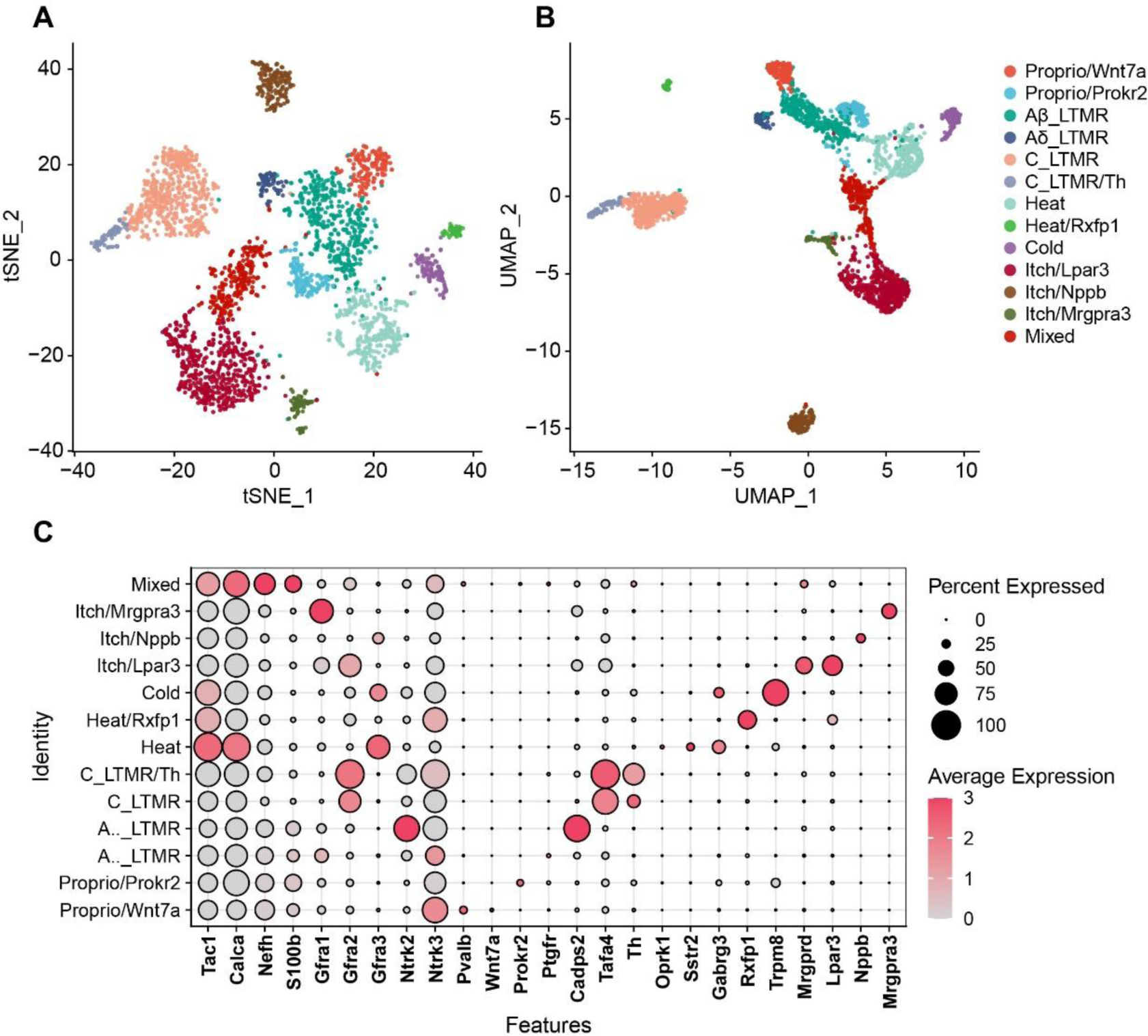
The heterogeneity of DRG neurons in mice with paclitaxel-induced CIPN. **A and B** t-SNE (A) and UMAP (B) plots of single cells profiled in the study, with each neuron type represented by a colored dot. **C** The dot plot of the neuron types in the DRG of paclitaxel-induced CIPN mice, marked by distinct genes.

### The Gene Ontology (GO) annotation of DEGs suggested potential alterations in nerve fiber and potassium-related currents in C_LTMR following paclitaxel treatment

We then performed the analysis of differentially expressed genes (DEGs) across all neuronal subclusters between Ctrl and Pac14 or Pac21. The gene expression in the minimum percentage of cells (min.pct) and the average log2 fold change (avg_logFC) between groups were set at 0.25. The results unveiled notable up- and down-regulated DEGs across all neuron types within the DRG at both day 14 and 21 post-paclitaxel injection (Figure 4A and B). Several genes, including Camk1d, RN18s, and Cdk8 were commonly up-regulated in multiple neuronal subtypes at two time points, indicating that these genes may be markers of paclitaxel exposure. Surprisingly, at the current sequencing depth, we were unable to quantified the expression of Atf3 gene (counts of all cell = 0), a marker of neuronal injury. It is worth noting that the C_LTMR and Itch/Lpar3 cluster had more DEGs compared to other clusters at both time points (Figure 4A and B), suggesting that these two clusters were most vulnerable to paclitaxel treatment at the transcriptome levels. In this study, subsequent analysis focused on C_LTMR, which has been associated with pain hypersensitivity in many models of chronic pain.

**Figure 4.**
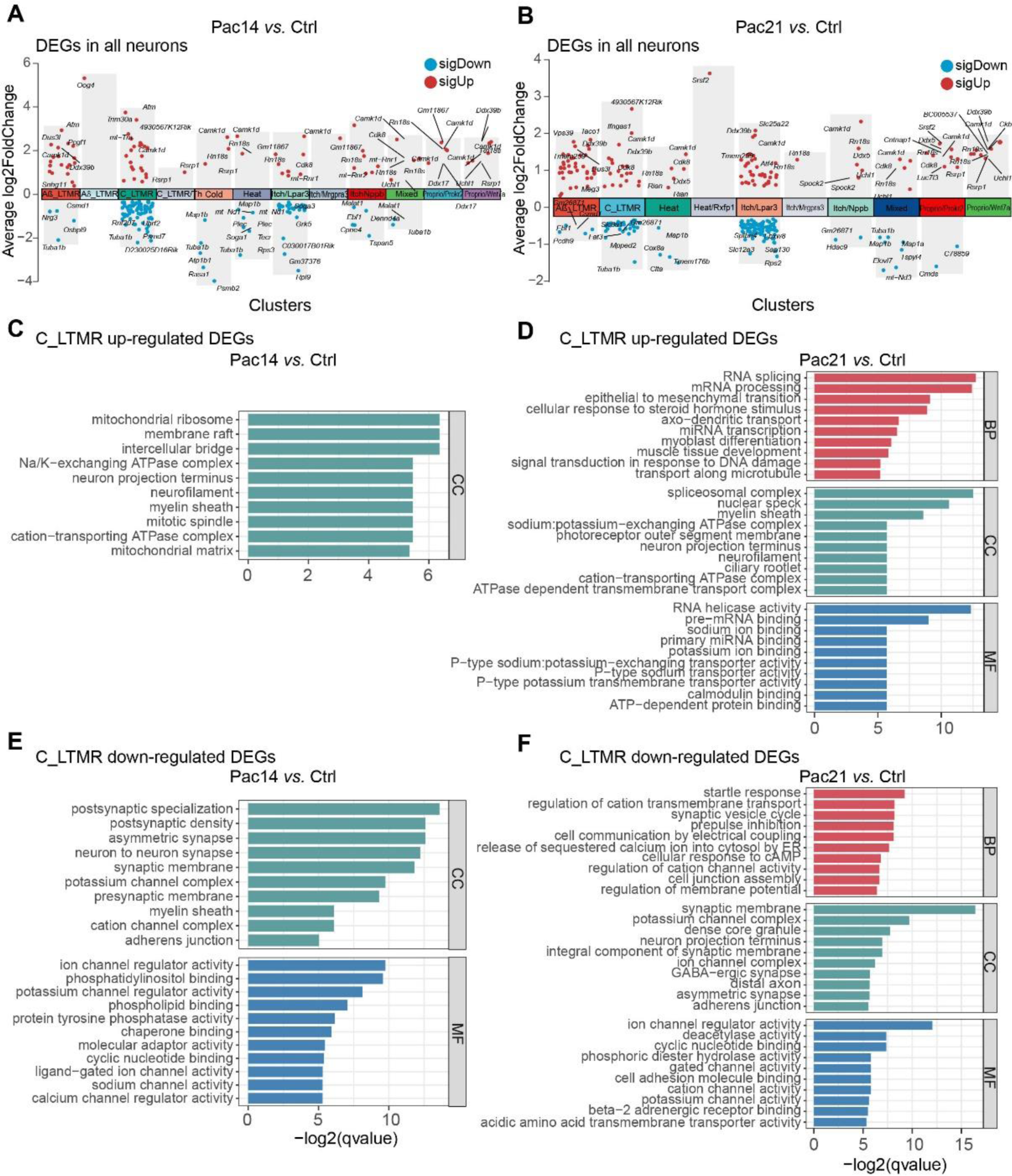
The transcripts regulated in all DRG neurons in mice with paclitaxel-induced CIPN. **A and B** The volcano plots showing the up- and down-regulated DEGs (differentially expressed genes) in all types of neurons in the DRG at both 14 (A) and 21 (B) days after paclitaxel injection. **C and D** The comparison of gene ontology (GO) enrichment of the up-regulated DEGs in C_LTMR neurons following paclitaxel injection at 14 (C) and 21 (D) days in mice. **E and F** The comparison of gene ontology (GO) enrichment of the down-regulated DEGs in C_LTMR neurons following paclitaxel injection at 14 (E) and 21 (F) days in mice. Top 10 significantly enriched GO terms, including biological process, cellular component, and molecular function, are shown.

Gene ontology (GO) analysis of the DEGs in the C_LTMR cluster was then performed. Analysis of up-regulated DEGs at day 14 post-paclitaxel injection failed to enrich biological processes (BP) and molecular function (MF) terms, but highlighted the association of cellular component (CC) terms such as “mitochondrial ribosome,” “mitochondrial matrix,” and “neuron projection terminus” (Figure 4C), suggesting that mitochondrion and axonal fibers of C_LTMR might have been affected. At day 21 post-injection (Figure 4D), enriched biological processes encompassed “RNA splicing,” “mRNA processing,” and “signal transduction in response to DNA damage” in C_LTMR neurons. Enriched cellular component terms included “spliceosomal complex”, “neuron projection terminus,” and “neurofilament,” while enriched molecular functions were linked to “RNA helicase activity” “pre-mRNA binding” and “calmodulin binding”. These results indicated that there may be persistent impacts on axonal fibers of C_LTMR and reorganization of RNA processing during paclitaxel recovery.

For down-regulated DEGs, enriched cellular component terms such as “postsynaptic specialization,” “postsynaptic density,” “neuron to neuron synapse,” and “potassium channel complex” were revealed at 14 days post-injection (Figure 4E). At 21 days post-injection (Figure 4F), enriched biological processes encompassed “synaptic vesicle cycle,” “cellular response to cAMP,” and “regulation of membrane potential” in C_LTMR neurons. Additionally, enriched cellular component terms included “synaptic membrane,” “potassium channel complex,” “GABAergic synapse,” and “asymmetric synapse,” while enriched molecular functions were associated with “ion channel regulator activity,” “deacetylase activity,” and “potassium channel activity.” This analysis implies that paclitaxel treatment could lead to dysregulation in potassium related currents, which might subsequently affect the excitability of C_LTMR.

### Gene Set Enrichment Analysis (GSEA) of transcriptomic changes in C_LTMR neurons

While enrichment analysis using DEGs reveals prominent treatment-related ontologies, it may overlook pathways associated with DEGs exhibiting subthreshold fold changes. To address this limitation, we employed the GSEA method to identify affected Gene Ontology Biological Processes (GOBPs) in C_LTMR cluster. The Seurat object of neurons was loaded to escape (Easy single cell analysis platform for enrichment) package and subjected to GOBP enrichment analysis. As depicted in Figure 5, aligning with the well-defined effects of paclitaxel treatment, the enrichment scores of several biological processes, including “microtubule-based transport” and “oxidative phosphorylation,” and “mitochondrion organization” were significantly decreased in C_LTMR neurons treated with paclitaxel for 14 days compared to saline (Figure 5A, B, C, and D). Notably, we observed that the enrichment scores of “Interleukin 17 production” were widely increased in multiple neuronal clusters after paclitaxel administration (Figure 5E). This elevation was particularly pronounced in various nociception and pruriception-related clusters, including C_LTMR, Cold and Itch/Lpar3, suggesting that Il17-related neuroinflammation might play a role in abnormal sensations after paclitaxel treatment.

**Figure 5.**
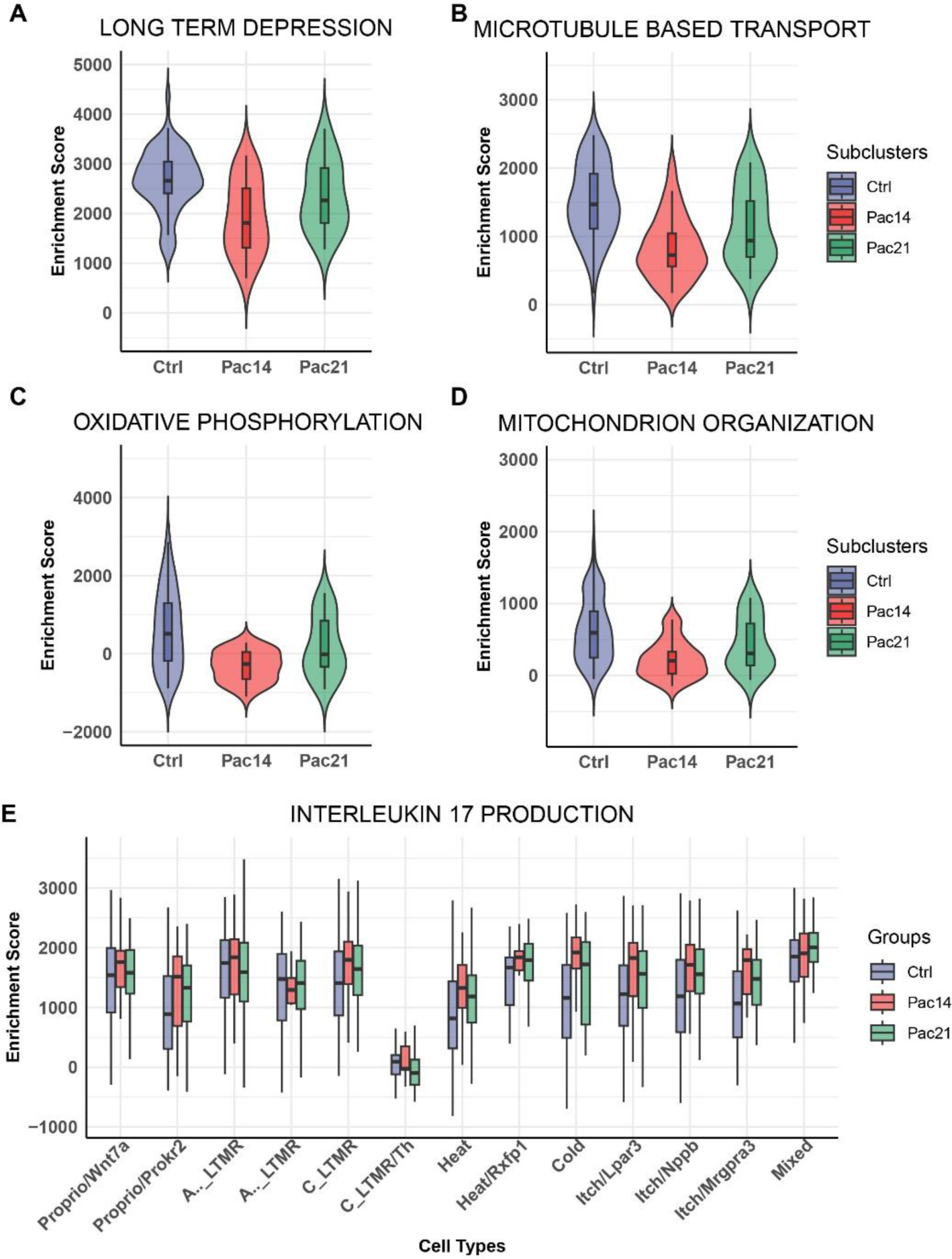
The gene set enrichment analysis of transcriptomic changes in C_LTMR neurons. **A, B, C and D** The desregulated terms in C_LTMR neurons in mice, showing that paclitaxel treatment was associated with several biological processes, such as “long-term depression” (A), “microtubule-based transport” (B), “oxidative phosphorylation” (C), and “mitochondrion organization” (D). **E** The enrichment score of Interleukin 17 production in C_LTMR neurons in mice after paclitaxel treatment (E).

### Pseudo-time analysis of C_LTMR cluster

Pseudo-time analysis enables the discovery of different cell states that exhibit subtle differences in cellular behaviours within a cell population. We subset the C_LTMR cluster and constructed pseudo-time trajectories to identify different cell states after paclitaxel treatment and genes with altered expression as the neurons transitioned between states (Figures 6A). A total of 9 states were identified across the trajectories (Figure 6B). Interestingly, the distribution of cell states varies among different groups, notably with a substantial increase in the prevalence of state 1 observed in two paclitaxel treatment groups. This result suggested that C_LTMR in state 1, positioned at the earliest point of the pseudo-time trajectory, represent a responsive state following paclitaxel exposure (Figure 6B-C). We then perform unsupervised clustering analysis of gene expression and categorize genes according to their expression trends across the pseudo-time frame. This assigned genes into 6 different clusters. Notably, genes in cluster 2, 4, and 6 exhibited a decreasing trend, whereas cluster 1 demonstrated an increasing pattern across the pseudo-time trajectory, indicating a close association with state 1 (Figure 6E). The GO annotations of clustered genes showed enrichment for various neurotransmission related terms, such as “synapse organization”, “axonogenesis”, “regulation of membrane potential” and “vesicle-mediated transport in synapse”. On the other hand, genes in clusters less likely associated with state 1 were more relevant to transcription modulation (e.g. “histone modification” and “chromatin “remodelling”). This result implied that the sustained presence of state 1 could directly contribute to abnormal pain processing after paclitaxel administration.

**Figure 6.**
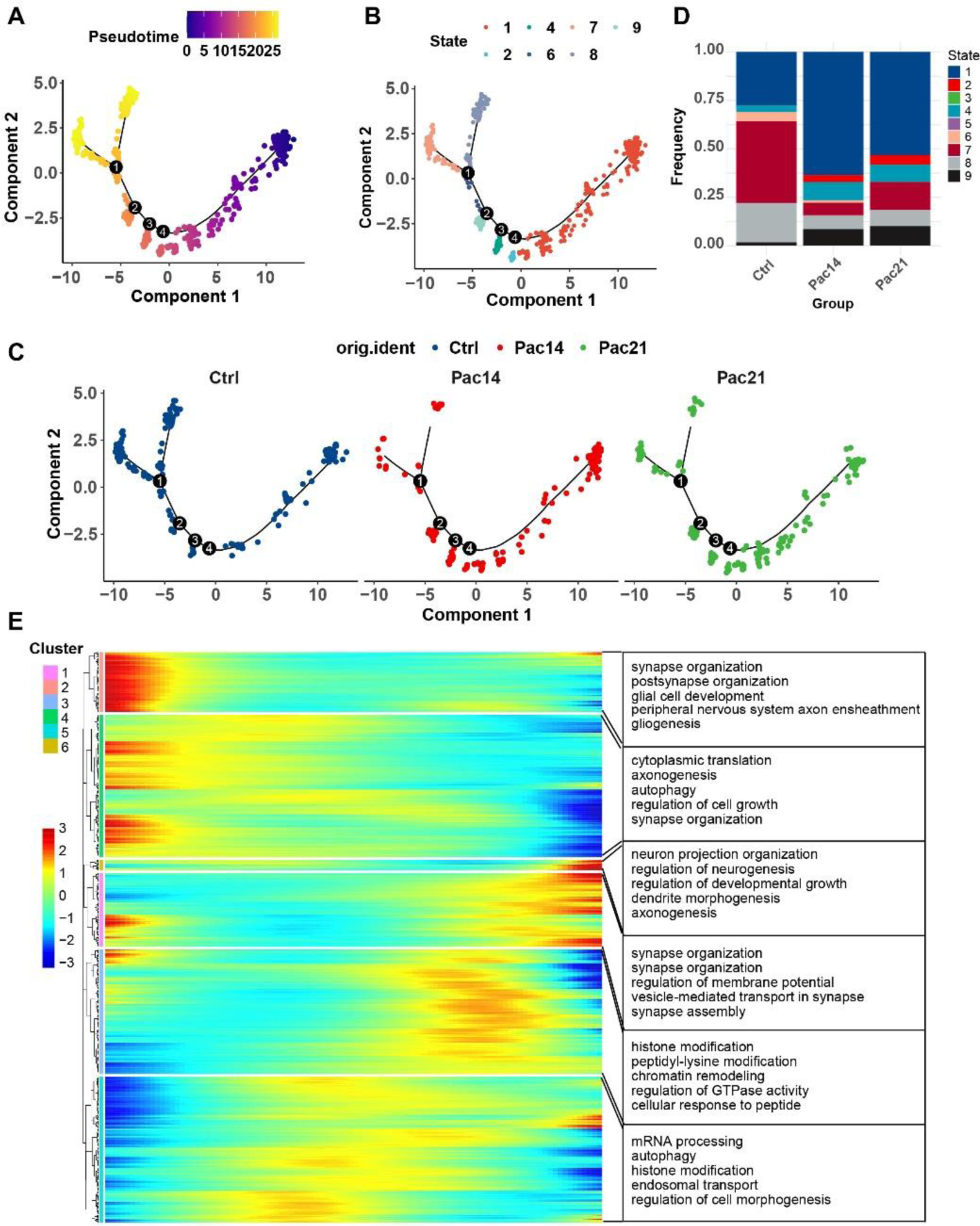
Pseudotime analysis of transcriptomic changes in C_LTMR neurons. **A** Pseudo-time trajectories show the transition of the process in paclitaxel-induced CIPN mouse DRG C_LTMR neurons. The dots represent cells, and the colors indicate neuron clusters or pseudotime. **B** Genes in C_LTMR neuronal switch process are classified into 9 modules based on their expressing patterns. **C** The separation of Modules 1, 2, and 3. **D** The bar graph showing the frequency ratio of each module. **E** Heatmaps that display the DEGs module based on their dynamic expression characteristics, as shown in pseudotime analysis, in C_LTMR neurons in paclitaxel-induced CIPN mice. Gene ontology (GO) enrichment analysis results were used to analyze the biological processes of each gene module.

### Gene interaction analysis unveiled gene modules associated with altered biological processes in state 1 C_LTMR

The products of genes interact physically or co-occur to accomplish biological processes. To identify gene modules potentially involved in altered biological processes in state 1 C_LTMR, we extracted the genes from cluster 2 (Figure 6C), which might have relevance to state 1, and constructed a gene interaction network using STRING (Figure 7A). The MCODE plugin of Cytoscape software revealed numerous densely connected subnetworks, notably highlighting subnetwork 1, which could explain changes related to mitochondrion (Figure 7B), and subnetwork 3, which could be relevant to alterations in neurotransmission (Figure 7D). Additionally, the existence of subnetwork 2 also suggested that paclitaxel might dysregulate the ubiquitination system in C_LTMR cluster (Figure 7C).

**Figure 7.**
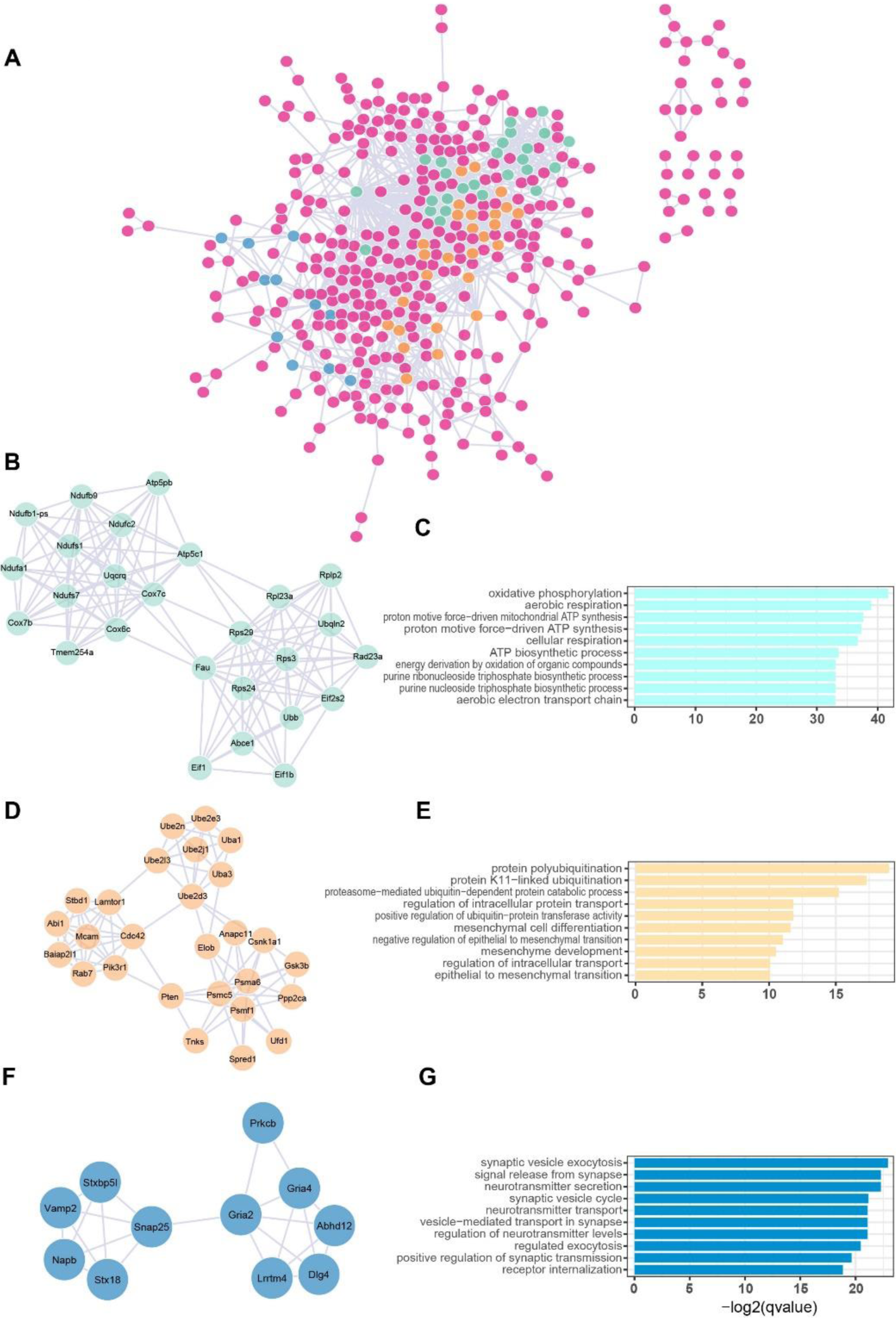
Gene interaction analysis in state 1 C_LTMR neurons in mice with paclitaxel-induced CIPN. **A** Gene interaction network analysis using STRING in state 1 C_LTMR neurons in mice with paclitaxel-induced CIPN. **B, D, and** F The MCODE plugin of Cytoscape software revealed numerous densely connected subnetworks, notably highlighting subnetwork 1 (B), subnetwork 2 (D), and subnetwork 3 (F). **C, E, and G** GO analyses were conducted to determine the biological processes associated with each of these subnetworks 1 (C), subnetwork 2 (E), and subnetwork 3 (G) in state 1 C_LTMR neurons in mice with paclitaxel-induced CIPN.

## Discussion

CIPN is a significant and often debilitating side effect of anti-cancer treatment, impacting patients’ ability to continue chemotherapy. Within 6-24 months after treatment, many patients experience neurotoxic symptoms, including numbness, stabbing and burning sensations, and pain in the extremities [22]. Therefore, it is crucial to find ways to prevent and treat CIPN without compromising the effectiveness of chemotherapy. This study sought to enhance our understanding of the mechanisms underlying CIPN by utilizing single-cell RNA sequencing in a mouse model with paclitaxel-induced CIPN. The sequencing revealed seven major cell types within the dorsal root ganglion (DRG), with neurons further categorized into 12 distinct subclusters using known markers. Notably, C_LTMR neurons exhibited the most significant transcriptomic changes following paclitaxel administration. Differential gene expression and gene ontology (GO) analysis highlighted suppressed potassium-related currents, microtubule transport, and mitochondrial functions in C_LTMR following paclitaxel treatment. Additionally, gene set enrichment analysis (GSEA) suggested an increase in Interleukin 17 production in C_LTMR after paclitaxel exposure. Pseudo-time analysis revealed nine distinct states (state 1 to 9) of C_LTMR, with State 1 showing a higher prevalence in paclitaxel-treated mice and altered neurotransmission properties, likely contributing to paclitaxel-induced pain hypersensitivity. This comprehensive exploration sheds light on the molecular mechanisms driving paclitaxel-induced neuropathic pain, offering potential avenues for therapeutic intervention. These insights represent a significant step forward in addressing the challenges posed by CIPN and its impact on cancer patients undergoing treatment.

Paclitaxel is a widely used chemotherapeutic drug in cancer treatment, but it can lead to peripheral neuropathy in a significant percentage of patients, with severe cases ranging from 22% to 100%. The development and severity of neuropathy depend on factors such as the dose, infusion duration, cumulative dose, and the presence of coexisting conditions like cancer and diabetes [23]. Neurotoxicity from paclitaxel is a common issue among chemotherapeutic agents, making the model of paclitaxel-induced CIPN a typical representation of CIPN in scientific research. This emphasizes the importance of gaining a deeper understanding of the mechanisms behind paclitaxel-induced CIPN and developing effective interventions to address this prevalent and impactful side effect of cancer treatment [23, 24].

At present, the exact mechanism of CIPN remains unclear, and most research, both domestically and internationally, has primarily focused on symptomatic treatment, with unsatisfactory results. Existing drugs for managing CIPN can weaken the anti-tumor effects of chemotherapy while providing only partial relief from pain. As of now, there are no ideal drugs or methods to effectively control or prevent CIPN, and the strategies in use are primarily symptomatic. In-depth research into the mechanism of CIPN is crucial to develop specific interventions and treatments based on the underlying pain generation mechanisms, rather than solely targeting symptomatic relief. In previous studies, we conducted transcriptome analyses of DRG in both oxaliplatin- and paclitaxel-induced CIPN, revealing substantial differences in the pathogenesis of these two CIPN models. Specifically, paclitaxel appeared to directly impact somatosensory neurons, while oxaliplatin showed a tendency to affect glial cells [13, 14]. Utilizing single-cell RNA sequencing, we aimed to further elucidate the cellular atlas of pain pathways and unveil cell-type-specific alterations. Our data revealed the identification of 12 distinct neuron subclusters within the DRG using known markers [15, 16]. Notably, C_LTMR neurons exhibited the most significant transcriptomic changes following paclitaxel administration. Previous reports have suggested that low-threshold mechanoafferents are involved in pain modulation [25] and inflammatory pain [26]. Our findings indicate that C_LTMR neurons in the DRG are primarily responsive to chemotherapy, which may correspond with the symptoms experienced by CIPN patients. This study sheds light on the specific cellular targets involved in paclitaxel-induced CIPN, providing potential insights for more targeted interventions and treatments.

The GO functional analysis revealed that the suppression of potassium-related currents, microtubule transport, and mitochondrial functions in C_LTMR neurons could be the primary mechanisms following paclitaxel treatment. These findings align with our existing knowledge of CIPN. Studies have reported that potassium channels play a role in the hyperexcitability of peripheral neuropathy [27, 28], and bortezomib has been shown to affect microtubule polymerization and axonal transport in rat DRG neurons [29]. U The ubiquitin proteasome system and microtubules are recognized as master regulators of central and peripheral nervous system axon degeneration [30]. Furthermore, mitochondrial damage and oxidative stress are well-documented events in CIPN [8, 31]. Additionally, gene set enrichment analysis (GSEA) indicated an increase in interleukin 17 production in C_LTMR neurons after exposure to paclitaxel. Research has shown that interleukin-17 regulates neuron-glial communications, synaptic transmission, and neuropathic pain following chemotherapy [32]. These findings provide valuable insights into the specific molecular pathways involved in paclitaxel-induced CIPN, potentially paving the way for targeted therapeutic interventions.

In summary, our study highlights the pivotal role of C_LTMR neurons in paclitaxel-induced CIPN. The suppression of potassium-related currents, microtubule transport, and mitochondrial functions in C_LTMR neurons as key mechanisms following paclitaxel treatment. These findings represent a significant addition to our understanding of CIPN pathogenesis and the identification of specific targets and signaling pathways within somatosensory neuron subtypes. Our research has the potential to contribute to the development of more targeted and effective approaches for treating CIPN by focusing on the specific cellular and molecular alterations associated with paclitaxel-induced CIPN.

## Declarations

### Ethics approval and consent to participate

All experiment procedures in this study were approved by the Ethics Committee of Animal Care and Use of the 6th Affiliated Hospital of Shenzhen University Medical School.

## Authors’ contributions

The authors’ contributions were as follows: X Liu and W Sun were responsible for the conceptualization and design of the study, data analysis, interpretation, and manuscript writing. W Sun, R Li, S Wu, and D Cao conducted animal experiments; R Li, X Zhang, S Wu, Y Jiang, Q Li, D Cao, D Xiong, and L Xiao reviewed the manuscript.

## Consent for publication

The paper has been read and approved by all authors. All authors approved the submission of this paper for publication. All authors confirmed that neither the manuscript submitted nor any part of it has been published or is being considered for publication elsewhere.

